# Dimension lifting in mental space for adaptive behavior in highly dynamic situations

**DOI:** 10.64898/2026.08.03.742413

**Authors:** Carlos Calvo Tapia, José Antonio Villacorta-Atienza, Gonzalo Aparicio-Rodríguez, Paloma Manubens, Sergio Díez-Hermano, Gerardo Oleaga, Valeri A. Makarov

## Abstract

Time compaction theory is a general framework explaining how a brain can efficiently deal with dynamic situations occurring in, e.g., sports games. It involves a geometric representation of the time dimension, which enables effective learning and strategic action planning. The theory has recently received experimental support in humans. However, its current computational model has an important limitation: it does not account for deliberate waiting and speed modulation, behaviors ubiquitous in natural environments. This work substantially extends the original model formulation by a dimensional lifting of an *n*-D workspace into (*n* + 1)-D mental space, where time remains geometrically embedded. The proposed biologically inspired computational model can generate adaptive behavior across increasingly complex situations, from navigation in everyday social environments to competitive sports. Furthermore, by actively conditioning the expected responses of other agents and stabilizing future predictions, we introduce the concept of uncertainty points in sequences of generalized cognitive maps to support the generation of adaptive strategies in interactive environments, where future prediction has a limited time horizon. Thus, we provide a mechanism for chaining short-term solutions into long-term strategies, which is illustrated by simulating the behavior of a player in a real football game.

**Author summary:** Humans often anticipate future interactions in dynamic environments. Many behaviors, such as avoiding other pedestrians, letting someone pass through a narrow corridor, or reproducing the kind of dribbling maneuvers performed by elite football players, require deciding not only where to move but also when to move. Existing theories suggest that the brain simplifies such situations by representing future interactions as static spatial maps, making them easier to learn and recall. However, current computational models cannot naturally account for common behaviors such as waiting, slowing down, or modulating speed. Here we show that these behaviors readily emerge if the model space is extended by an additional virtual coordinate that encodes accumulated waiting rather than physical time. The proposed model simultaneously admits a wide variety of behaviors, including speed modulation, multigoal decisions, and compound actions, while preserving the principles of time compaction. We illustrate the model in everyday situations and by reproducing two real football plays, comparing the observed behaviors with model simulations. Our results suggest computational principles through which the human brain may efficiently represent, memorize, and exploit dynamic situations.

## 1 Introduction

Humans and animals constantly interact with a time-evolving environment and other living beings. Therefore, their nervous systems have developed neuronal substrates and cognitive mechanisms allowing them to anticipate the future and adapt their behavior accordingly. At the ultimate level, the ability to represent or mentally model the environment has emerged, allowing for strategic planning and the performance of complex motor actions such as navigation, escape, or pursuit. The study of these mechanisms started from the initial hypotheses proposed by Darwin [1]. Then, Tolman provided the first experimental evidence [2], whereas nowadays spatial cognition has been accepted as a key component of human experience [3].

Spatial cognition relies on internal representations of the environment [4], supported by distinct neuron populations: place cells fire at specific locations [5], grid cells fire at the vertices of a hexagonal grid internally mapped onto space [6], object-vector cells encode distances and orientations relative to objects [7], and head direction cells signal head orientation [8]. In static environments, these specialized neurons, along with others, contribute to the generation of the cognitive map [2, 5, 9, 10], an internal model of the surrounding space stored in memory that enables effective decision making [11], flexible behaviors [12], and the generalization of information to new experiences [13].

Time-evolving environments can be represented internally via a generalization of standard cognitive maps [14, 15]. The underlying mechanism, termed time compaction, encodes, within a purely spatial map, the locations where the subject or *agent* could potentially interact with other moving and static elements, including other living beings. Thus, the time dimension is embedded in space, and the resulting *generalized cognitive map* (GCM) can be readily stored in memory and recalled for future situations, as it happens with standard cognitive maps. Therefore, GCMs support flexible behaviors and real-time interactions in dynamic situations and serve as a basis for cognition, influencing higher cognitive functions [16, 17]. Time compaction is implicated in learning and decision-making [18] and modulates memory [19]. The soundness of this cognitive mechanism is supported by its presence in various mammals. Humans [18] and rats [20] have been experimentally shown to use time compaction, and similar results have been observed in bats [21].

The existing mathematical GCM formulation implicitly assumes that adaptive behavior is achieved through spatial route selection, while the agent’s movement speed remains fixed [14]. Such a reduction restricts the application of GCM to many relevant behaviors, such as fighting, pursuing, or fleeing, which critically depend on the timing of actions rather than solely on the selection of spatial routes [22–24]. Different forms of adaptation ubiquitous in natural environments involve deliberate waiting, acceleration, or the strategic timing of actions [25–27]. Current computational models cannot encode them and hence remain incomplete.

The conceptualization of functional approaches to understanding how nervous systems implement flexible executive behaviors in dynamic situations is novel in computational biology. Recently, it has been suggested that the brain may use the concept of the blessing of dimensionality in high-dimensional spaces [28, 29] to represent and manipulate complex concepts [30]. Such dimension lifting has also been exploited in the context of artificial agent navigation to enhance adaptive behavior. The concept of state-time space extends the traditional state space or workspace approach by incorporating time as an additional dimension [31–33]. It allows for speed variations and can enable safe and flexible navigation in dynamic environments [34–37]. However, this approach achieves flexibility by explicitly representing time, which is suboptimal for compact representations [33]. Moreover, a natural speed limitation cannot be readily achieved, which may lead to unfeasible solutions. Therefore, the challenge is not simply to represent speed modulation, but to do so while preserving the principles of time compaction.

In humans and many other animals, motivation plays a central role in decision-making [38, 39]. Given its significance in daily life, it would be inefficient for the brain to adjust its cognitive processes to a specific motivation and then reevaluate everything if that motivation changes. Therefore, it is beneficial for cognitive mechanisms to be capable of simultaneously generating multiple potential actions and goal-oriented decisions [40]. As a result, incorporating the multigoal property into the GCM model presents another challenge.

In this work, we extend the original GCM model by adding a virtual dimension to the workspace. The extra spatial dimension has a much more evolved nature and represents cumulative “waiting” time. This dimensional lifting preserves the principles of time compaction while enhancing adaptive behaviors and context-sensitive decision-making. In particular, it allows the agent to change speed on purpose and perform complex maneuvers with multiple goals simultaneously encoded within a single GCM. The proposed computational model is illustrated across scenarios of increasing complexity, including situations taken from real football games. Rather than reproducing specific motor commands or optimizing trajectories, we show how dynamic real-world situations can be represented in mental space to support cognition, adaptive behavior, and strategic planning of motor actions.

## 2 Dimension lifting in mental space

### 2.1 Wait dimension

Let us consider a typical problem of navigation in a social environment: a man (*agent*) walks to a door while a woman crosses his path (Fig. 1A). Although the workspace is three-dimensional (3-D), the navigation problem effectively develops on a 2-D plane. We can project all elements of the situation onto the (*x, y*)-floor, which represents the effective workspace, which we use below.

**Fig 1.**
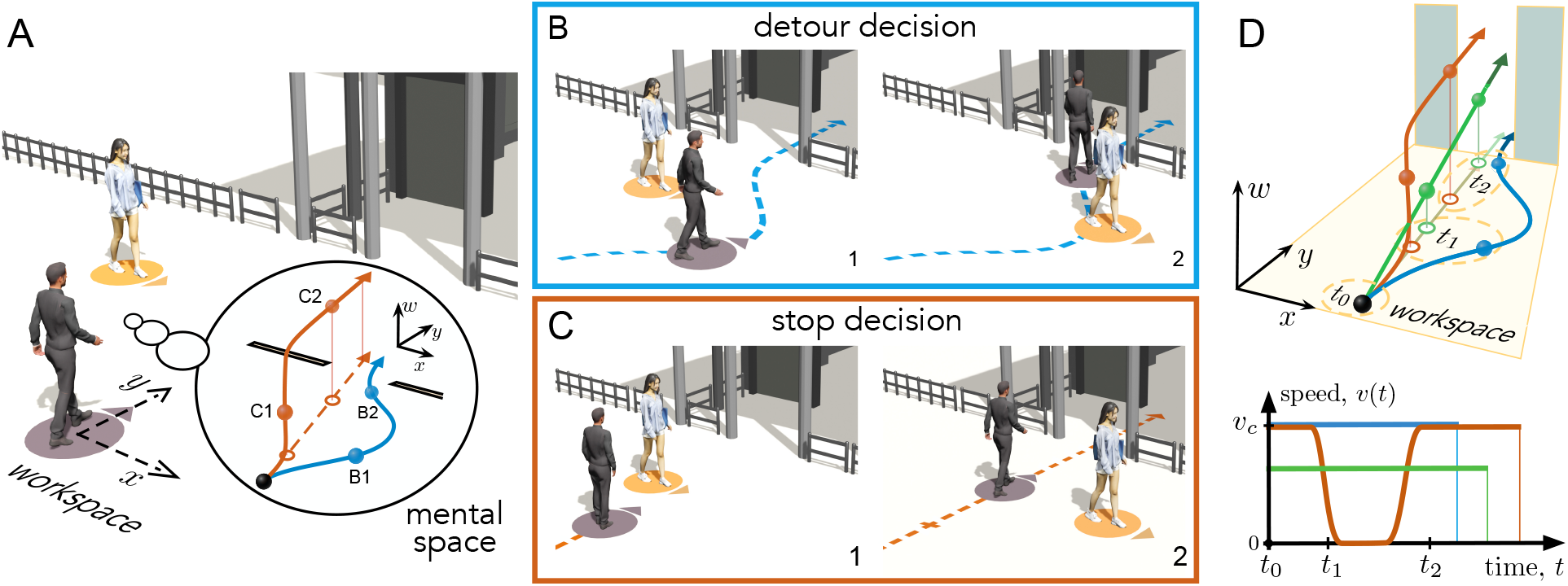
The concept of mental dimension lifting. **A**. A typical dynamic situation: a man approaches a building entrance while a woman crosses his path. The workspace is effectively 2-D (the (*x, y*)-plane). The man can either 1) walk without stopping and cross in front of the woman, or 2) stop, let her go first, and continue walking. These decisions are represented in the extended 3-D mental space (the bubble inset) by curves starting from the man’s initial position (black dot). The blue curve corresponds to the non-stop or detour solution with constant speed; the wait dimension does not change, *w*(*t*) = 0. The red curve shows the stop solution. While the man does not move, the wait coordinate grows (i.e., *dw/dt >* 0), thus compensating for the deceleration in the 2-D workspace. The projections of the decisions onto the (*x, y*)-workspace depict the corresponding pathways in physical space (the blue curve and its projection coincide). **B**. Execution of the detour decision in the workspace (dashed curves). Two time instants correspond to the blue dots in panel A. **C**. Same as in B but for the stop decision. In panel 1, the man stands still while the woman passes by. **D**. Different decisions in the mental (*x, y, w*)-space produce different trajectories in the workspace. The additional green curve corresponds to a decision to move with constant speed *v < v*_c_. The pathway is the same as in red. The agent’s speed depends on the dynamics along the *w*-coordinate. Note that the agent’s position at the same time instants *t*_1_ and *t*_2_ depends on the adopted decision. The green curve may cause a collision with the woman and hence is invalid and must be discarded by a mental model.

To avoid a collision and get through the door, the man (the agent) has two options: either stop and let the woman pass, or detour around her. Although the detour decision is longer than the stop one, it could be shorter in time, since the agent does not stop or slow down. Thus, we have a typical multiobjective optimization problem. Both solutions can be optimal in different scenarios.

Following the time-compaction framework, we aim to embed different decisions in a geometric space, so the time dimension plays no role. To achieve this, the original time-compaction paradigm assumes a constant agent’s speed (other objects can have time-varying speeds) [14]. We relax this restriction and assume that the agent can vary his speed within an interval *v*_a_ ∈[0, *v*_c_], where *v*_c_ is the maximum or comfortable speed. Time compaction is then achieved through dimension lifting, which occurs within the agent’s mental space.

The bubble inset in Fig. 1A shows the 2-D problem in the *mental* 3-D space. The mental space is intended to capture an internal cognitive representation of the environment and the agent’s potential interactions with it. It includes the third *virtual* dimension *w*, which we call *the wait dimension*. It extends the (*x, y*)-workspace and enables encoding of changes in the agent’s speed. In the mental (*x, y, w*)-space, all trajectories are purely geometric objects and have the same meaning, and can be compared easily. The movements along any 3-D trajectory occur at constant speed *v*_c_. To obtain a path in the 2-D workspace, we project the corresponding 3-D trajectory onto the (*x, y*)-plane.

The detour trajectory (blue curve in the bubble inset in Fig. 1A) maintains the maximal speed in the workspace. Thus, no wait occurs, and the wait coordinate does not change, *w*(*t*) = 0, i.e., the mental trajectory lies in the *w* = 0-plane. The stop decision (red curve) includes a period during which the agent does not move in the workspace. To compensate for this behavior, the red curve rises in the *w*-direction in the mental space while preserving its projection on the ground, so the agent’s position in the (*x, y*)-floor does not change, but the accumulated wait is evident. Therefore, the virtual *w*-dimension encodes the agent’s accumulated waiting along its pathway in the real workspace.

Importantly, the wait dimension should not be interpreted as physical time. Two points with identical *w*-values may correspond to different time instants, since the elapsed time depends not only on the accumulated waiting but also on the spatial path traversed. Rather, *w* quantifies the amount of waiting the agent accumulates over a decision. Decisions lying on the plane *w* = 0 coincide with their projections on the (*x, y*)-plane and therefore do not involve any deceleration, as illustrated by the blue trajectory in Fig. 1A.

Figures 1B, C depict the executions of the red and blue decisions in the workspace. While along the blue pathway (Fig. 1B, time instants 1 and 2), the agent makes a detour without stopping, Fig. 1C1 shows the agent standing still, waiting for the woman to pass by (see the corresponding red dot in Fig. 1A). We also note that the *w*-dimension does not encode speed itself. Instead, speed modulation emerges naturally as a geometric consequence of traversing trajectories embedded in the 3-D mental space.

The virtual dimension provides flexibility in decision-making beyond the dichotomy between standby and comfort-speed navigation. To isolate the effect of the wait dimension, we, for the moment, ignore the presence of the woman in the scene and construct an additional third decision in the mental space (Fig. 1D, green curve). This decision shares the same projection to the workspace with the red curve, and hence the agent also moves along the straight line. However, its speed is constant, lower than in the detour case (blue curve), *v < v*_c_. Then, the agent’s decision is 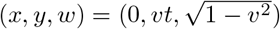. We now observe that at time instant *t*_1_ the projection of the green trajectory onto the (*x, y*)-workspace overtakes the red one. At time *t*_2_ the agent’s positions over the blue and green decisions are similar in the workspace, but the *w* values are significantly different. Thus, the man arrives at the door at the same time but over different paths. However, remembering that the woman crosses the agent’s path, we observe that the green decision may cause a collision, which invalidates such a trajectory, and this decision must be discarded. We also observe that the agent taking different trajectories arrives at the door at different time instants (Fig. 1D, speed panel).

The considered example illustrates how the wait dimension enriches the repertoire of decisions available to the agent. All decisions are purely geometric paths. Thus, in some sense, we reduce the time-evolving problem to a static one. Then, we need a computational model to construct all possible paths in the mental space and discard those that are not valid due to collisions with other entities in the environment (see below). The described mechanism is general and supports multiple decisions in any situation, regardless of complexity. Thus, the dimension-lifted mental space provides a substrate for an efficient internal representation of all potential decisions.

### 2.2 Computational model of mental space

From a biological perspective, decision-making relies on the internal exploration of alternative courses of action and the evaluation of their potential consequences [41]. To implement this paradigm, we use a neural medium through which a nonlinear wave of excitation propagates and interacts with other activity, thereby simulating the presence of other objects. It allows: 1) representing the exploration of the agent’s potential decisions in the 3-D mental space, and 2) evaluating the interactions between these decisions and the predicted evolution of the environment. Then, valid decisions become geometric objects that can be explored and compared within the same representational space.

Earlier, the exploration dynamics has been implemented using neuronal lattices [14], cellular automata [42], and the dynamic fast marching method [43]. Here we adopt a continuous three-dimensional formulation for analytical and computational convenience. Such an approach is typical for modeling the macroscopic behavior of large neural networks using coarse-grained variables (see e.g. [44]).

For convenience, we formulate the model in dimensionless variables. The reference length is chosen as the personal-space radius of the agent *ρ* = 1. Thus, the agent is represented by the unit disk. All distances are therefore expressed relative to this characteristic length. Likewise, the reference time is defined through the propagation speed of the excitation wave, which corresponds to the agent’s maximum comfortable speed, yielding *v*_c_ = 1.

Let **x** = (*x, y, w*) ∈^*T*^ ℳ ⊂ ℝ^3^ and *t* ∈ [0, *T*] be the coordinate vector in the mental space ℳ and the physical time. *T >* 0 is the look-ahead time horizon for mental prediction. The neural excitation field *u*(*t*, **x**) ∈ℝ governs the exploration process, and its space-time evolution is described by the following reaction-diffusion partial differential equation:

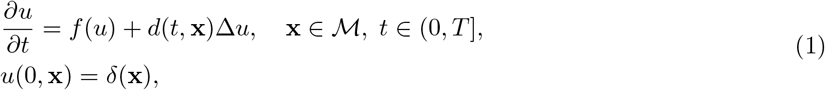

where 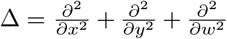 denotes the 3-D Laplacian operator. The local dynamics of the neural medium is determined by the cubic reaction term:

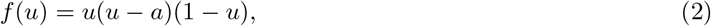

where *a* ∈ (0, 1*/*2) is the activation threshold of the neural tissue. The local dynamics defined by *f* (*u*) is bistable. There are two attractors, *u* = 0 and *u* = 1, whose basins are separated by the unstable equilibrium *u* = *a*. Thus, we consider regions with *u*(*t*, **x**) *< a* as being in a quiescent (rest) state, while regions with *u*(*t*, **x**) ≥ *a* correspond to activated neuronal populations. The initiation of the exploration of ℳ starts from the agent’s current state corresponding to a point-source excitation centered at the origin of the egocentric system. In these settings, the mental space ℳ is bounded by a ball of radius *Tv*_c_ = *T*, which is essential for implementation.

In a free, unobstructed isotropic domain where the diffusion coefficient is constant (*d*(*t*, **x**) = *d*_0_), Eq. (1) admits a sigmoid-like traveling wavefront,

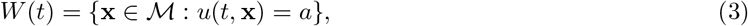

which expands spherically at an asymptotically constant velocity depending on the diffusion coefficient *d*_0_. This expanding wavefront simultaneously explores a continuum of all possible decisions that radiate from the initial position (for details, see [43]).

The presence of dynamic opponents and static boundaries in the workspace (e.g., the woman and walls in Fig. 2A) alters the diffusion topography. Such entities occupy space and hence disturb wavefront propagation. To take them into account, we represent external objects by occupied regions *b*_*j*_(*t*) ⊂ ℝ^2^, which may be either static or dynamic depending on the predicted evolution of the environment. Humans are modeled as disks whose radii *r*_*j*_ correspond to their personal spaces. For example, the region occupied by the woman is given by:

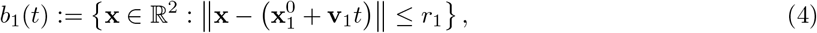

where 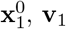, and *r*_1_ denote her initial position, velocity, and personal-space radius, respectively. More generally, *b*_*j*_(*t*) represents any region of arbitrary shape occupied at time *t*. Then, the complete set of external objects is:

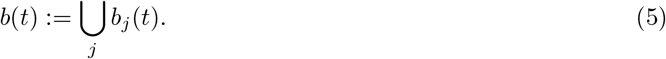

**Fig 2.**
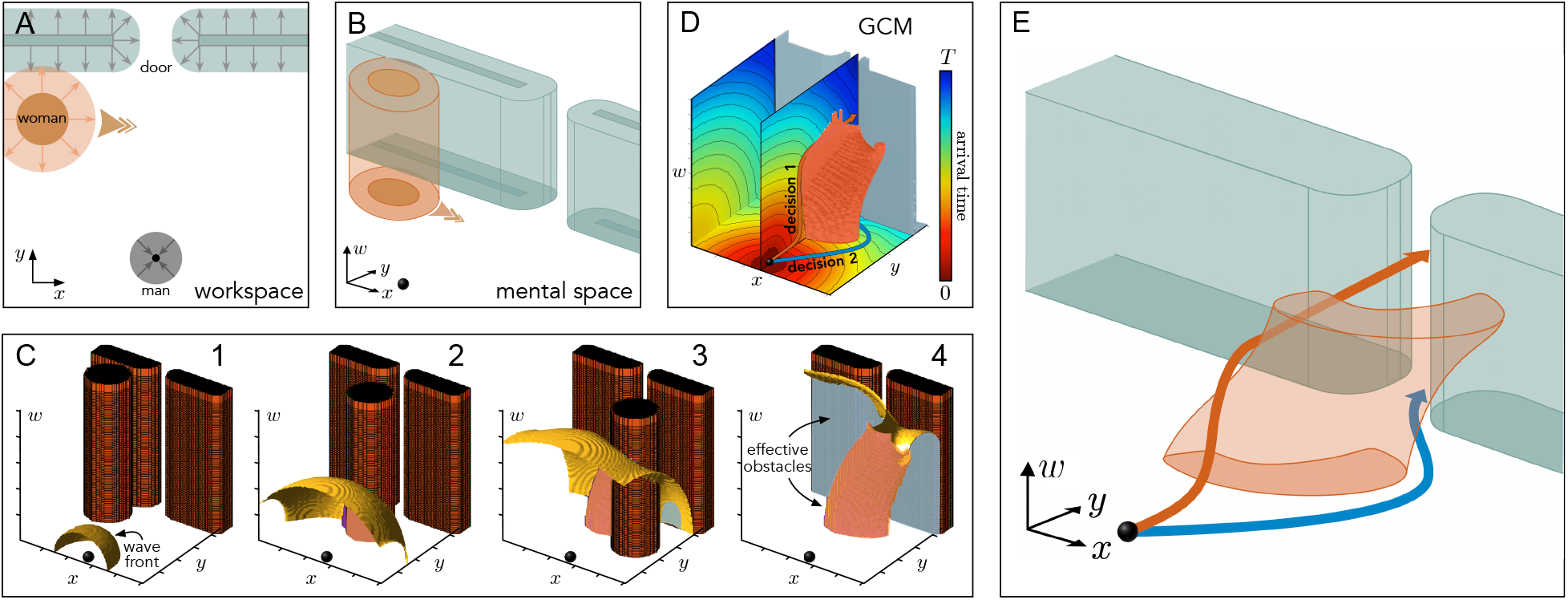
Computation of potential decisions in the mental space for the situation described in Fig. 1A. **A**. The situation in the 2D workspace. Blue, red, and gray figures represent the walls, woman, and man, respectively. For mental simulation, the man is reduced to a single point, while the external elements are enlarged accordingly (arrows of corresponding colors, Eq. (6)). **B**. Dimension lifting in the mental space. Objects in the 2-D workspace are extended along the *w*-axis (Eq. (7)), while the agent is a point at the origin (black dot). **C**. The wavefront (yellow) propagates at speed *v*_c_ and explores the mental space (simulation of Eq. (1)). When the wavefront reaches the brown objects (representing the woman’s future states and the walls), the effective objects (reddish and gray shapes) are progressively generated (Eq. (8)). **D**. Generalized cognitive map formed after time compaction. The GCM contains a static field describing the arrival time of the wavefront (Eq. (10)). Multiple decisions can be obtained by ascending the gradient field. **E**. Sketch of the decision-making in the mental space. The reddish figure represents possible collisions between the man and woman. The agent cannot traverse through. Any decision curve starting from the initial position (black dot), ascending in the time field *F* (**x**), and avoiding effective obstacles is a valid collision-free solution (two of them are shown by blue and red curves, see also Fig. 1). The workspace is a square 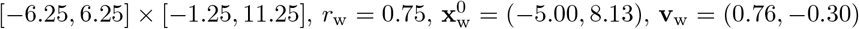.

To account for the finite size of the agent, we adopt a configuration-space representation in which the agent is collapsed to a point while surrounding objects are enlarged accordingly (Fig. 2A):

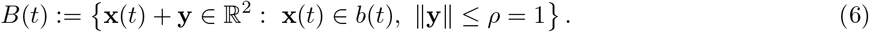

Then, the agent’s movement is described by a 1-D curve.

Given the region *B*(*t*) occupied by the enlarged environmental objects in the workspace (i.e., in the plane *w* = 0) at time *t*, we can represent it in the mental space ℳ . Since collisions are independent of the virtual wait coordinate *w*, the occupied region undergoes a spatial extrusion along this virtual dimension, giving rise to the set of extended objects (Fig. 2B):

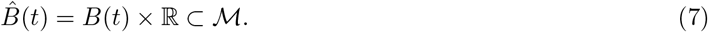

When the expanding wavefront *W* (*t*) intersects the boundary of an extended object, 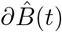, in the mental space, it instantly creates an impenetrable region in ℳ. This boundary interaction is formally integrated into the system by defining the dynamically growing set of effective objects (Fig. 2C):

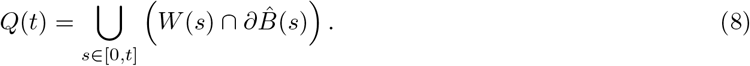

Consequently, the binary diffusion coefficient *d*(*t*, **x**) drops abruptly to zero within the blocked territories, rendering them strictly impenetrable to the wavefront:

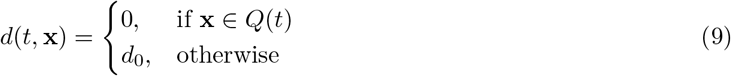

As the wavefront diffracts and slides around the effective objects, it progressively maps out all kinematically feasible decisions. Once the wavefront has fully explored the reachable domain *A {* **x** ∈ ℳ : *u*(*T*, **x**) ≥ *a*, the stationary GCM is obtained by extracting the potential field of arrival times (Fig. 2D):

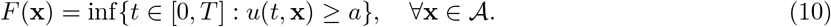

The potential field *F* (**x**) mathematically defines the GCM by capturing the minimum arrival times of the neural propagation across the mental space. Decisions can then be extracted by ascending through this field (e.g., blue and red curves in Figs. 2D, E), yielding collision-free trajectories adapted to the predicted evolution of the environment.

Figure 2E shows a simplified cartoon of the condensed spatiotemporal structure of the situation, reduced to a static representation. In this representation, the physical interpretation of the dimensions reveals the model’s core concept: while the coordinates (*x, y*) define the actual geometric path in the workspace, *w* introduces a crucial degree of freedom in the allocation of time. Rather than directly prescribing a kinematic velocity, the lifting into the *w*-dimension allows a single physical location (*x, y*) to be evaluated across multiple arrival windows. Consequently, the mental space allows the agent to decide not only where to position itself, but also precisely when to do so.

### 2.3 Numerical implementation

To solve Eqs. (1)-(10) numerically, we use the Strang splitting method [45], a second-order accurate member of the family of Fractional Step Methods [46]. Briefly, the dynamics (1) is decomposed into reaction and diffusion parts:

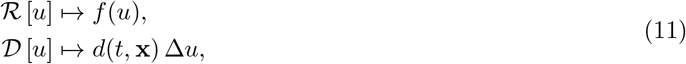

which are treated independently using their own numerical schemes.

The mental space ℳ was discretized on a regular Cartesian grid *{***x**_*ijk*_*}*, where **x**_*ijk*_ = (*x*_*i*_, *y*_*j*_, *w*_*k*_)^*T*^, and time was discretized as *t*_*n*_ = *n*Δ*t*. Then, 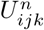 denotes the numerical approximation of the neural activity *u*(*t*_*n*_, **x**_*ijk*_). The binary diffusion coefficient *d*(*t*, **x**) was updated at each time step according to Eq. (9). To simplify notation, its spatial and temporal dependence is omitted in the numerical expressions below.

For the reaction step, we apply a second-order Runge-Kutta method (also called Heun’s method):

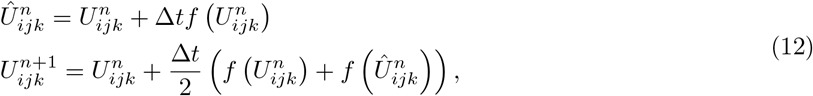

where the nonlinearity is evaluated componentwise: 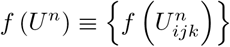. Thus, the reaction step is handled explicitly through the discrete operator:

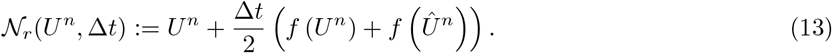

The diffusion term is discretized using the standard finite-difference scheme:

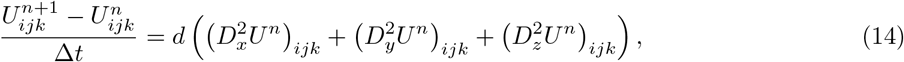

where the second-order finite-difference operators are defined along each spatial direction. For example, for the *x*-direction we have:

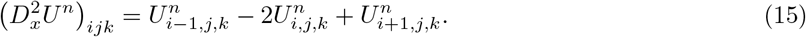

Thus, we get:

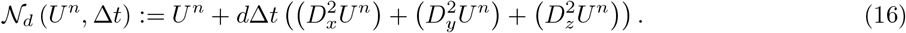

The complete Strang scheme can now be formulated. Given the discrete solution 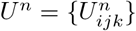 at time *t*_*n*_, the following intermediate states are computed:

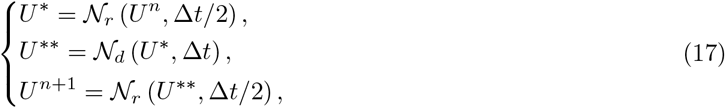

where *N*_*r*_ and *N*_*d*_ denote the numerical operators associated with the reaction and diffusion steps, given by (13) and (16), respectively.

## 3 Results

In Sect. 2.2, we observed that the mental dimension lifting provides flexible decision making in a simple but typical situation (Fig. 2). Let us now study other human behaviors of increasing complexity.

### 3.1 Single GCM admits multiple goal-motivated behaviors

As discussed in the Introduction, motivation plays a central role in decision-making. In the example developed in Figs. 1 and 2, the agent considered different decisions but with the same goal: to reach the door. However, GCM can also generate decisions whose projections onto the workspace yield trajectories leading to distinct destinations, thereby linking distinct behavioral outcomes to different motivations. Moreover, to construct a GCM, it is not necessary to assign the final point for the agent, i.e., GCMs are goal-free.

To illustrate this, we consider a daily situation where people move in a narrow space. Figure 3A shows a woman walking in a cafeteria and a man approaching her with a tray. In this scenario, the woman (agent) may want to give way to the man either by keeping moving forward or staying in place (top and bottom panels, respectively). Although her goal varies, both cases are described within the same GCM. Figure 3B shows the GCM built in the mental space and the projection of two solutions into the workspace.

**Fig 3.**
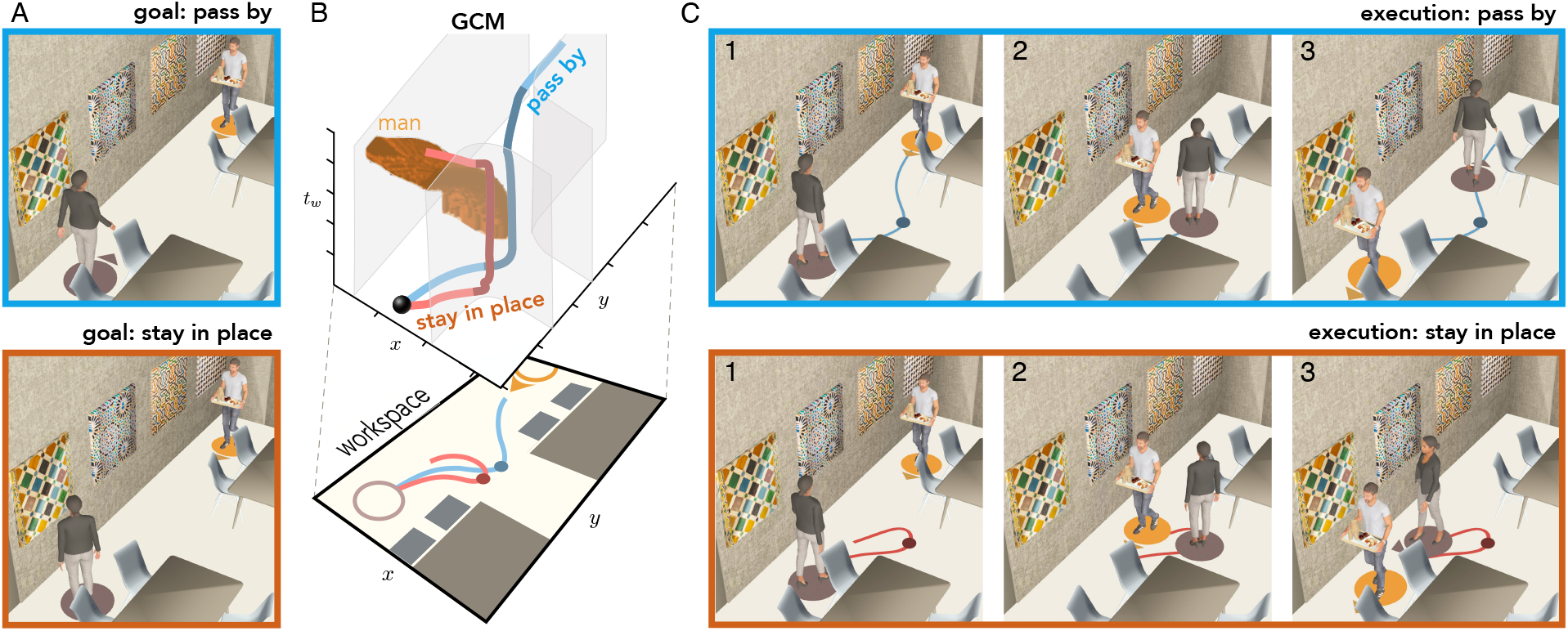
Decision-making under diverse motivations using a single generalized cognitive map. **A**. A woman (agent) in a cafeteria wants either to move forward (top), avoiding a man approaching with a tray, or to stare at the painting on the wall (bottom). **B**. The GCM encoding both decisions (for visual clarity, only the effective object representing the man is shown). According to her motivation, the woman extracts the appropriate decision (blue or red curves) to move in the workspace. Both solutions allow the man to pass, but the blue continues forward, whereas the red returns to the initial position. Both decision curves rise along the *w*-axis, which implies that the woman stops between the tables (blue and red dots in the workspace). **C**. Consecutive snapshots of the execution of the decisions.

The resulting decisions lead to markedly different behaviors: one yields a trajectory that continues along the narrow passage (blue curve), whereas the other returns the woman to her initial position (red curve). Figure 3C illustrates the two intelligent and realistic behaviors, similar to those observed in everyday situations.

This example illustrates a fundamental consequence of introducing the wait dimension. The additional degree of freedom not only increases the versatility of the decisions contained in a GCM (studied in Fig. 2), but also enlarges the set of situations for which feasible solutions exist. Both behaviors observed in Fig. 3 rely on the possibility of temporarily waiting while the man traverses the narrow passage. Importantly, different behaviors do not arise from a change in the environment, which remains identical in both cases, but from a change in the agent’s goal. The same GCM therefore supports multiple adaptive decision-making without requiring new mental simulations.

### 3.2 GCM provides decision-making in highly competitive sports

Figure 4A shows a highly dynamic situation occurring in football. A player (agent, A) aims to score a goal while three opponents try to avoid it. Using the observed movement of these players, the agent’s mind represents the entire play through a single GCM containing multiple feasible decisions. The player then executes the best goal-driven decision.

**Fig 4.**
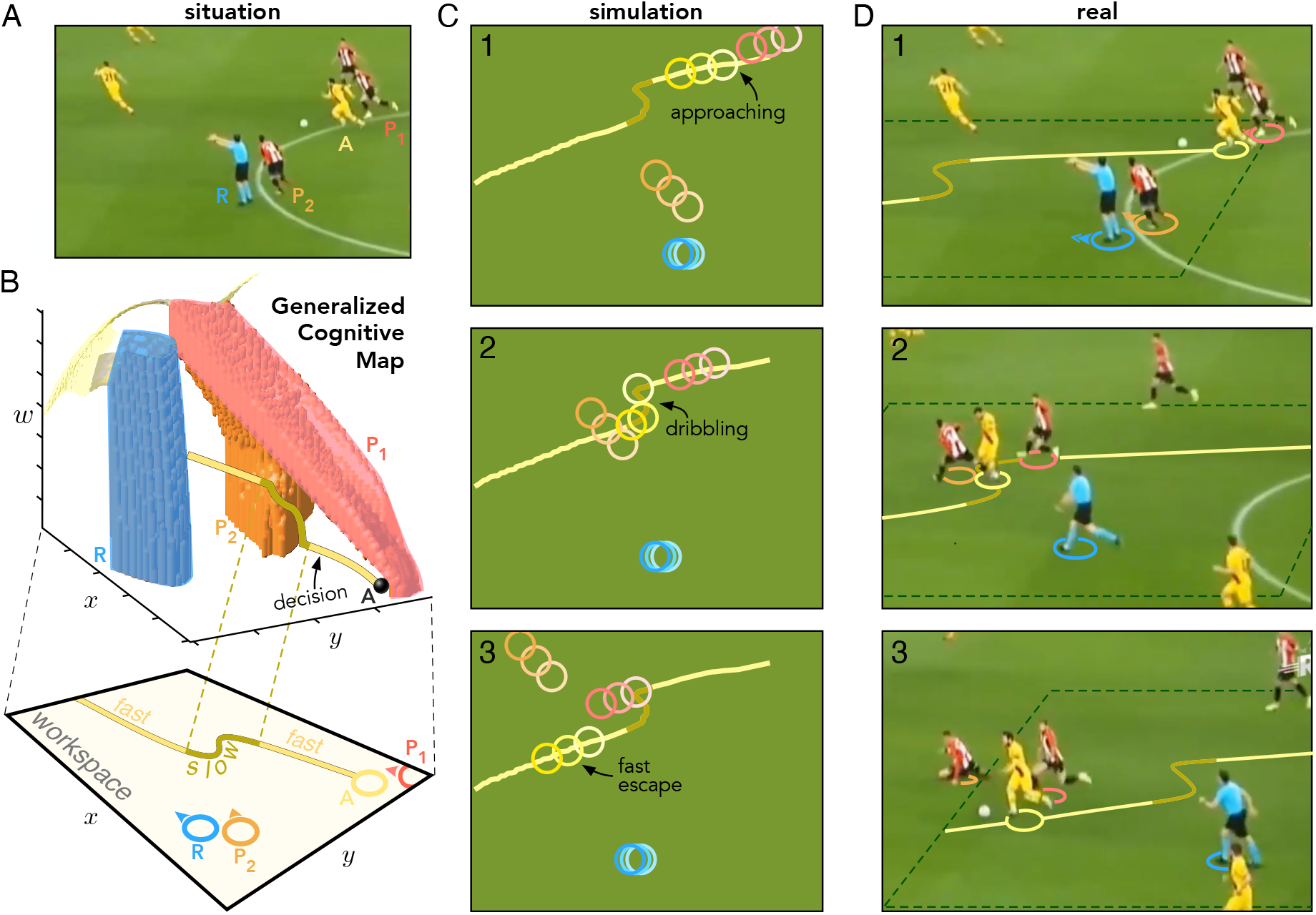
GCM models the behavior of a football player. **A**. A player (agent, A, yellow) drives the ball while avoiding rivals (red and white striped, marked as P_1_ and P_2_). **B**. GCM built for the situation (top 3-D image). Around the agent’s initial position (black dot), three effective obstacles emerge (red, orange, and blue shapes), corresponding to the two rivals (P_1_ and P_2_) chasing the agent and the referee (R). The GCM yields a solution (yellow curve) whose projection onto the workspace (2-D bottom plane) allows the agent to: 1) escape from player P_1_ (initial fast segment), 2) dribble past player P_2_ (slow segment), and 3) face the goal (final fast segment). Note that the dribbling involves a speed change provided by the elevation of the decision curve along the *w*-axis (dark yellow segment). **C**. Three consecutive snapshots of the execution of the simulated solution. The color code is the same as in B; superimposed circles with increasing color intensity correspond to progressively increasing time instants. **D**. Three consecutive snapshots of the real execution performed by the agent.

We modeled the situation by constructing a 2-D workspace and projecting all players involved in the scene, as well as the referee (Fig. 4B, workspace). We also estimated their initial velocities to build the time-dependent set of extended objects used in simulations. Then, we computed the corresponding GCM (Fig. 4B, top). As a target, we set a location near the football goals, which yields a decision involving a dribble (yellow curve). Such a decision causes a surprise effect on the opponents (players P_1_ and P_2_) and allows the agent to avoid them in the mental space.

The workspace projection of the dribbling decision (Fig. 4B, bottom) leads to the simulated execution on the playing field shown in Fig. 4C. This decision is qualitatively similar to the actual decision the agent makes to resolve the situation (snapshots in Fig. 4D). The key to success in this decision is the abrupt change of the speed and direction of running during the dribble, which confuses the opponents (dark yellow segment in Figs. 4B, C, and D). Thus, the GCM approach allows us to simulate decision-making in highly competitive sports.

### 3.3 Long-term strategic planning by concatenating GCMs

In Sect. 3.2, we have shown that GCMs enable decision-making with abrupt speed changes, which favors the surprise effect on opponents and scoring a goal. However, in a more complex situation, it may lead to a collateral problem: the appearance of uncertainty points, as the reliability of subsequent predictions decreases. Thus, uncertainty about the predictions on which the agent acts can invalidate long-term decision-making.

A solution to mitigate the emerging uncertainty is to concatenate possible decisions into a chain, thus creating an action strategy [47]. A biological mechanism supporting such a concatenation uses memory. Since GCMs are static sets (time is embedded in space), they are readily memorized during training and can be further retrieved to cope with highly dynamic situations in real time.

Figure 5A shows a player (agent, marked by A) surrounded by three opponents (P_1_-P_3_) whom he is unlikely to dodge with a single dribble, as it happened in the situation shown in Fig. 4. Performing a similar mental processing as in the previous example, in a first dribble the agent again leaves behind two opponents (Fig. 5A, panels 1-2). However, the dribble now leads to an uncertainty point, as the last defender likely stands in his way towards the football goals (Figs. 5A, panels 3-4). Thus, a new decision must be concatenated with the previous one.

**Fig 5.**
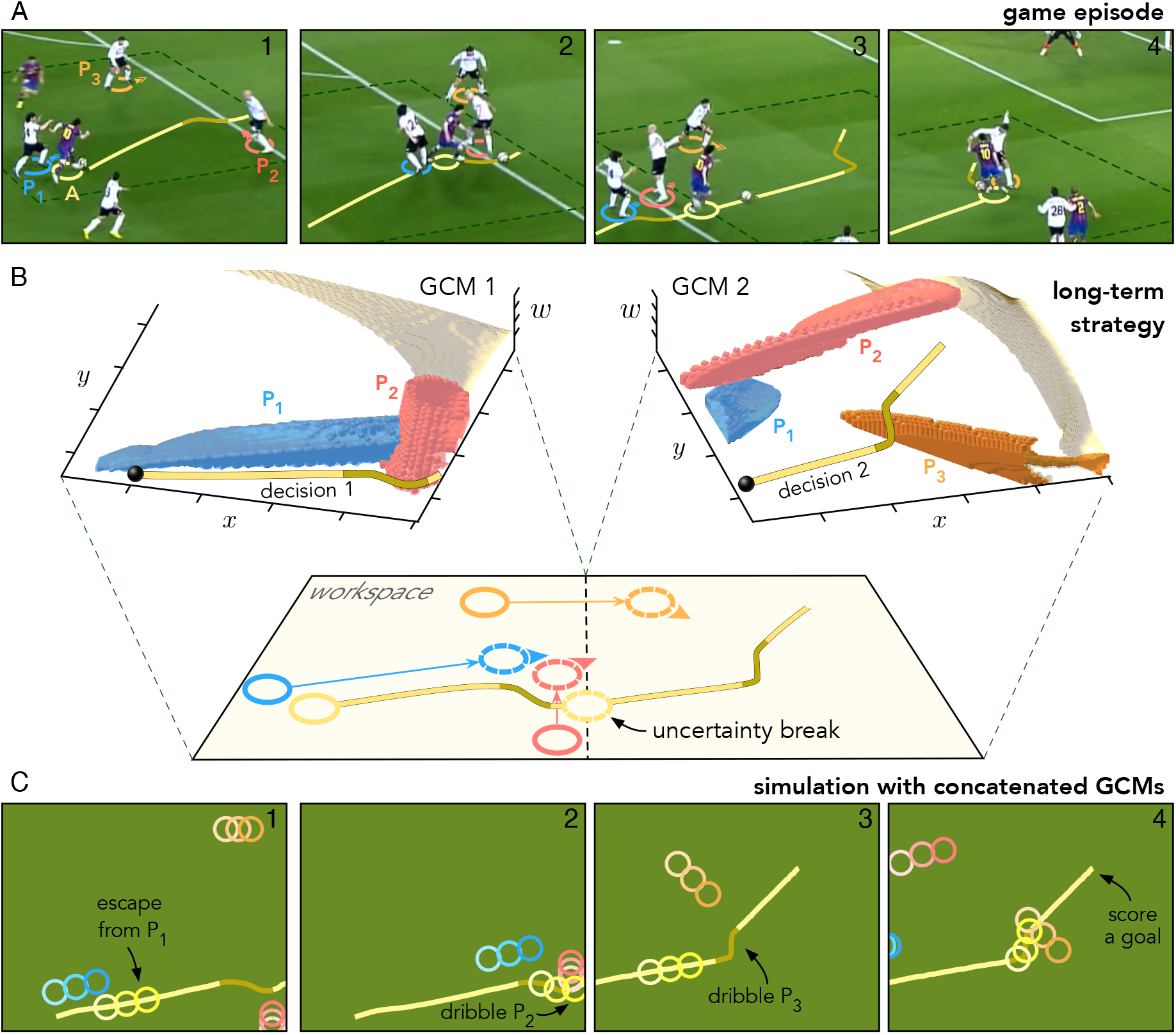
Building a long-term strategy by concatenating generalized cognitive maps. **A**. Consecutive snapshots of a real football play. A player (agent, marked by A in a yellow circle) escapes from three rivals marked by P_1_-P_3_ (blue, red, and orange circles) by dribbling twice (dark yellow segments). The agent first escapes from rival P_1_ (blue circle, snapshot 1), then dribbles to pass rival P_2_ (red circle, snapshot 2), next enters the area (snapshot 3), and finally dribbles to pass rival P_3_ (orange circle, snapshot 4) and scores a goal. **B**. Building long-term concatenated strategy. GCM 1 (top-left 3D image) solves the problem with players P_1_ and P_2_ (blue and red shapes). The projection of the decision (yellow curve) to the left half of the workspace shows the escape from player P_1_ and the dribble to rival P_2_ (game episode shown in panels A 1–2). Then, an uncertainty point arises, since the initial actions of the rivals change, which breaks the first solution. The new positions and directions of the opponents and of the agent himself (dashed circles and arrows in the workspace) constitute the input to obtain the second GCM (top-right 3-D image). A new effective obstacle corresponding to the player P3 appears (orange shape), inducing the second dribble (dark yellow segment). **C**. Consecutive snapshots of the play execution from the simulated GCMs. Snapshots 1-2 and 3-4 correspond to the left and right halves in panel B, respectively. Superimposed circles with increasing color intensity correspond to progressively increasing time instants.

Using the same methodology as in Sect. 3.2, we built the first GCM that allows us to reproduce the decision made by the agent in the first dribble (Fig. 5B, left). Its projection to the workspace yields the agent’s trajectory shown in the left half of Fig. 5B. Then, the final arrangement of the opponents and the agent location from the first map establish the initial conditions for generating the second GCM and performing the second decision (Fig. 5B, right). To improve the similarity with the real game episode, we corrected the intermediate positions by using panel 3 in Fig. 5A. The long-term mental planning is achieved by concatenating two consecutive GCMs, separated by the uncertainty point (Fig. 5B, dashed circles).

Assuming that the predictions of the time evolution of external parties are valid, concatenating the solutions chosen in each GCM yields a global decision that reproduces the one taken by the agent (compare Figs. 5 A and C). This shows the plausibility of long-term strategic planning based on GCMs for highly dynamic real-world situations with uncertainties.

## 4 Conclusions

In this work, we have developed a functional approach to understanding how the brain implements long-term motor behaviors in highly competitive human scenarios, abundant in sports games. Based on time compaction theory, we have proposed a novel concept of dimension lifting that leverages the so-called waiting coordinate *w*, extending an *n*-D workspace to the (*n* + 1)-D mental space. To describe the mental space dynamics, we have introduced a biologically motivated computational model describing the mean-field excitation in a neuronal tissue. A spherical wave propagating in the neuronal tissue constructs a generalized cognitive map of a situation that contains different decisions, which an agent can further execute. Importantly, GCMs are static sets that are ready to be memorized and then retrieved from memory for building complex real-time behaviors.

Our numerical studies showed that the proposed dimension lifting extends the representational capacity of the plane *n*-D formulation. Previous models may often lead to freezing behavior in crowd situations despite the presence of obvious solutions. The new mental space encodes not only alternative spatial routes but also temporally structured actions involving speed modulation and waiting behaviors. The model capacity is based on the natural assumption that the agent has a maximum or comfortable speed *v*_c_. Then, slopes of a decision curve in the *w*-dimension indirectly induce changes in the agent’s speed. Note also that different decision curves sharing the same projection to the workspace correspond to running over the same pathway but with different velocity profiles. Thus, dimension-lifted GCMs explain a great variety of behaviors observed in humans.

We have illustrated the model capacity in different dynamic situations of increasing complexity. First, we observed that the model can successfully cope with typical daily situations. It provides different decisions to reach the same goal by optimizing either the execution time or the traveled distance. Moreover, a single GCM admits multiple goal-driven behaviors. At the same time, the agent can either decide to stay in place or walk forward. Both decisions imply specific movements provided by the GCM to avoid collisions.

Second, we simulated behaviors of a high-rank player in football games. In one game episode, the player was able to surprise and escape with a ball from three opponents. The model GCM constructed from the initial situation provided a trajectory (path and speed modulation) qualitatively identical to that used by the player. Then, we modeled an even more complex scenario that includes uncertainty in decision-making. The player’s complex behavior in one episode produces high uncertainty in subsequent predictions. Long-term planning then requires concatenating different decisions in a chain. We have shown that the GCM approach is well suited to achieve the objective. The simulated behavior composed from two GCMs was qualitatively similar to that observed in the game.

Thus, the proposed biophysically motivated computational model is capable of constructing internal representations of highly dynamic and competitive situations in the form of generalized cognitive maps. The obtained GCMs are behaviorally flexible, multiobjective, and enable concatenation for long-term strategic planning. They tightly reproduce real human behaviors.

## 5 Discussion

A conceptual description of human behavior is relatively new in computational biology. Recent advances in computer science have pushed large language models to levels that were incredible for futurologists and AI experts only a decade ago [48]. At the same time, progress in modeling complex human motor behavior is less impressive. One reason may be the computational complexity associated with intellectual and motor activity supported by the presence of 1.6 × 10^10^ and 6.9 × 10^10^ neurons in the cerebral cortex and cerebellum, respectively [49].

The proposed computational model describes functional mechanisms behind cognitive decision-making for motor behaviors. Earlier models exhibited certain limitations when applied to highly competitive human scenarios, such as sports games. The dimension lifting substantially enlarges the representational capacity of generalized cognitive maps by converting temporal aspects of behavior into geometric properties of the mental space. The waiting coordinate differs significantly from the straightforward inclusion of the time coordinate proposed before [32]. It naturally includes a restriction on the maximum speed the agent can develop and makes the mental space homogeneous, hence preserving the principles of time compaction. Thus, it simplifies the process of building internal representations, which may be advantageous from an evolutionary point of view.

Two main lines of experimental evidence support the biological foundation of the model. First, time compaction has been observed as a cognitive mechanism in humans [18] and rats [50], and it is compatible with observations in bats [21]. Second, the brain is known to encode elapsed time between events [51], with continuous input-output flow mechanisms proposed to support this capacity [52]. The GCM-based model incorporates waiting as an additional dimension within the internal representational space, which is explored in parallel with the workspace, in line with these paradigms. GCMs minimize the encoded information by focusing on the spatial mapping of possible future interactions that contain a biologically plausible set of decisions to cope with time-changing situations [53]. The biological plausibility of the GCM has been illustrated by examining dynamic scenarios of increasing complexity, from everyday contexts to real football plays. Our work shows that human decisions correspond to solutions generated by the model, suggesting that GCMs may capture computational mechanisms supporting learning and the generation of adaptive decisions.

Memory is essential for decision-making during navigation [54]. It involves cognitive maps as internal representations of the environment [55]. Concurrently, growing evidence points toward compositional memory, according to which complex memories are constructed from simpler ones [56]. Specifically, in static environments, cognitive maps emerge from the composition of simpler maps [13]. Time compaction allows for the memorization of complex dynamic situations based on the composition of simple static representations. Thus, we extend the compositional paradigm to more realistic scenarios, while also enabling real-time decision-making. Previous computational studies have demonstrated the potential of GCM as a cognitive framework for complex mental processes, such as memorizing dynamic situations [17] and the compositional generation of strategies in real time [47].

More broadly, the proposed functional framework is compatible with several influential perspectives in contemporary cognitive neuroscience. By representing dynamic situations in terms of their potential future interactions and associated decisions, GCM aligns with ecological views emphasizing action-relevant information and affordances [57, 58]. At the same time, it provides an explicit mechanism for their internalization and storage [57, 59, 60]. Likewise, the compression of complex interactions into compact predictive structures is consistent with event segmentation theories [61–64], which posit that experience is organized into discrete cognitive units, and with predictive processing frameworks [65–68], in which cognition relies on the generation and updating of predictions. From this perspective, GCM may provide a computational substrate through which dynamic experiences are compressed, memorized, and exploited to support adaptive decision-making and strategy generation.

These strategies consist of a mental sequence of two or more concatenated decisions, explored in this study within the context of navigation. The functional basis of the GCM is the combination of predicting the behavior of external dynamic elements with the mental simulation of the subject’s own potential decisions. In this sense, in real dynamic situations involving other cognitive agents, predicting their behavior depends on what the subject might do and vice versa; this recursivity is critical for strategy generation [15].

Specifically, we propose that the mental generation of a strategy is based on the concatenation of GCMs: the decision derived from each of them remains valid up to a certain point of uncertainty. These points are defined as the agent’s position within each GCM where their decisions would cause changes in the other actors that would subsequently alter the original prediction. Ultimately, the core advantage of our model lies in its information-processing efficiency, as it optimally resolves complex strategies through a sequence of simple, static representations of a dynamic environment that would otherwise be hardly manageable.

This aligns with the soccer context analyzed here, which is an appropriate example of a complex dynamic environment in which experience is key to success. Players require training to deal with complex plays. They do not need a novel generation of strategies for each scenario they face, but rely on previous experience consolidated through the performance of similar plays. Time compaction provides the appropriate functional framework for this. By embedding the temporal dimension into space, GCM-based encoding optimizes learning and memorization of complex dynamic situations. Furthermore, representing experiences as static points in a multidimensional space allows the definition of appropriate metrics for establishing relationships between such situations in terms of hierarchy, similarity, and causality [17].

Potential-field methods usually resort to gradient descent techniques or their modifications to obtain trajectories [69]. Our approach does not restrict traversal to strictly following the gradient; it suffices to causally follow the field values. At this juncture, we distinguish between safe trajectories avoiding collisions with obstacles and feasible trajectories, those that the agent can effectively execute given its mechanical constraints. The fact that two points can be connected by ascending the field indicates the existence of a safe transition between them. However, proximate field values could correspond to locations that are sufficiently distant in workspace to demand unattainable velocities or accelerations. This discrepancy highlights an inherent limitation of the model: the safety of a trajectory does not guarantee its feasibility.

The framework used addresses fundamental questions in cognitive neuroscience: how the brain understands complex, dynamic environments to generate effective and efficient decisions, and how it builds long-term strategies. Our results suggest that dimension-lifted GCMs support both capabilities. The approach enables the generation of potential decisions adapted to different motivations by incorporating speed changes, stops, and alternative trajectories, thereby capturing behavioral features essential for efficient adaptive behavior. On this basis, strategies emerge through the concatenation of yet-unexecuted decisions, internally explored across successive GCMs and connected by uncertainty points that delimit the predictive horizon of each representation. Importantly, the decisions and strategies generated by the model qualitatively reproduce key features of real football plays, linking the proposed computational mechanisms to observed human behavior in highly dynamic competitive scenarios.

